# Pathway Volcano: An interactive tool for pathway guided visualization of differential expression data

**DOI:** 10.1101/2024.11.21.624662

**Authors:** Thomas M OConnell

## Abstract

**Summary:** The increasing size and complexity of omics datasets can make effective visualization and interpretation very challenging. Differential expression datasets as from RNA-sequencing and proteomics can contain many thousands of features and even after filtering for significance there can be hundreds or thousands of features to consider. A common tool for visualizing this type of data is the volcano plot where each feature is plotted as log2 transformed fold change along the x-axis and the negative log10 of a p-value along the y-axis. These plots provide a useful visualization of the largest and most significant changes, but the majority of the features are often crowded and overlapping near the “cone” of the volcano. In order to provide a biologically informative way to simplify the visualization and interpretation of these types of datasets we developed the Pathway Volcano tool. This R-Shiny based software utilizes the Reactome API to select specific pathways and then filters the volcano plots to show only the data associated with those pathways. In this manner, many of the significant features in the crowded section of the volcano plot can be revealed to support the impact of specified pathways. This tool provides a range of interactive features to interrogate the data along with the ability to download png files and tables with pathway associated data.

**Availability and implementation:** Pathway Volcano is a freely available R Shiny package. The program was developed in R version 4.3.3 using R Studio version 2024.09.1 Build 394. Running the app locally requires the packages ggplot2 (1), plotly (2), shiny (3) dplyr (4) and ReactomeContentServer (5). The full code and documentation including example datasets are available at https://github.com/thoconne/PathwayVolcano

## 1. Introduction

The application of omics technologies in biomedical sciences is becoming increasingly prevalent. Among the most widely used approaches are RNA-sequencing and proteomics. In both cases, the differential expression of either genes or proteins are measured to understand the biological differences between groups or conditions (6-8). With increasing advancements in the analytical technologies, the datasets resulting from these studies are becoming increasingly large (9). Many tools are available for the statistical analysis of these datasets which typically yield list of differential expression including fold changes and an assessment of significance including DESeq (10) and MAGenTA (11) for genomics data along with MaxQuant/Perseus (12) and MSstats (13) for proteomics.

A common visualization approach used in the interpretation of differential expression data is the volcano plot. This type of plot typically presents the log2 transformed fold change along the x-axis and the negative log10 of an adjusted p-value along the y-axis. Typical datasets from RNA sequencing analyses can yield well in excess of 10,000 transcripts. After significance testing, datasets are reduced, but can still retain hundreds or even thousands of significant features. Thus, volcano plots, even when presenting only the significantly altered transcripts are extremely crowded making the interpretation of the data very challenging. Often only the metabolites that are extremely significant receive any attention while the majority of significant features go uninterpreted.

A number of specialized tools have been developed to generate high quality volcano plots for differential expression data. Mullan *et al*., developed an R Shiny application called ggVolcanoR which allows for highly customizable volcano plots for use with RNA sequencing and proteomics data (14). Different sets of analytes can be colored and labeled by virtue of significance and directionality of change. The EnhancedVolcano tool is an R package available on Bioconductor which provides a similar set of volcano plot customizations but currently requires experience with R programming in order to generate these plots (15). Even with the ability to highlight analytes with varying magnitudes of fold change and significance, a challenge remains that many statistically significant expressions found near the very crowded “cone” of the volcano are being ignored.

To overcome this challenge and simplify the data visualization a different approach to filtering and highlighting the data is required. As the ultimate goal of many omics investigations is the identification of perturbations to specific biological pathways, we developed a method to filter and visualize the data based on biological pathways. This is accomplished by matching the experimental data against the lists of genes and proteins in the Reactome Database using the Reactome API (16). The resulting volcano plots are filtered to contain only the entities in the experimental dataset that are found in a specifically queried pathway. In this manner, the biological pathways suggested by some of the most significant changes can be visualized to potentially reveal additional changes in the more crowded region of the plot that support this change. Hypotheses regarding other pathways that are suggested from additional types of analyses e.g. over-representation analysis in Reactome (17) or GSEA (gene set enrichment analyses), (18) can also be used to guide the examination of specific pathways.

## 2. Results

The Pathway Volcano tool was written in the R programming language using the R-Shiny framework along with the Plotly package for interactivity (2,3). The Reactome database API is called using the ReactomeContentService4R package (5). The program can be easily loaded and launched from R-Studio without any additional programming and can be easily hosted on routine server hardware.

### 2.1 Data upload

Figure 1A shows the Data Input Panel of the tool which allows the user to upload a CSV file with differential expression data. The file can have any number of columns, but there are three required columns labeled (1) GeneSymbol (2) log2FoldChange and (3) padj. For simplicity the gene symbols are coerced to the mammalian (non-human) format with a capital first letter and subsequent lower case letters. When using proteomics data, the associated gene name can be used and the data upload is identical.

**Figure 1.**
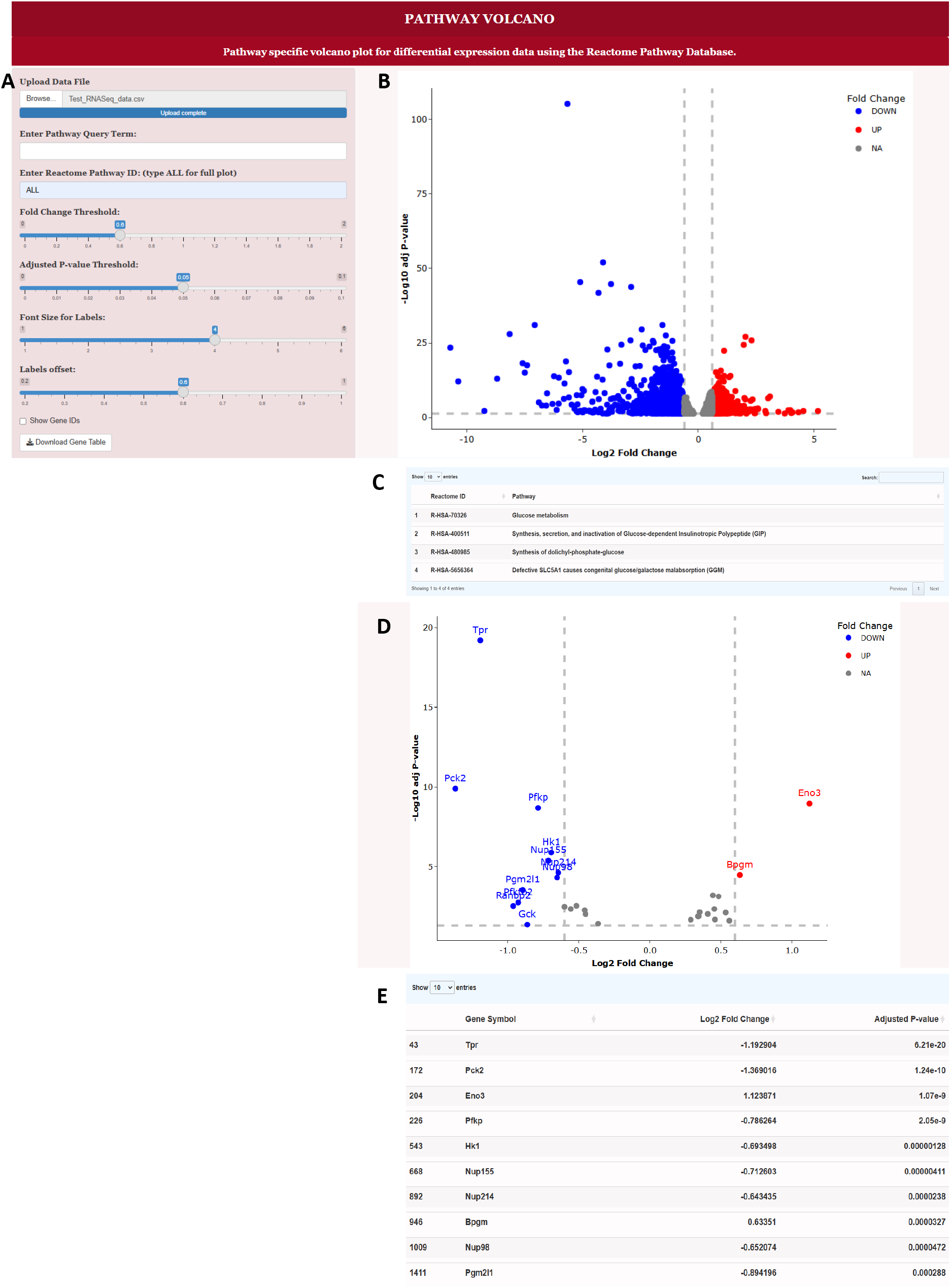
Interface to the Pathway Volcano Tool and example output (A) Data upload and query panel. In this panel, the user can upload the experimental data, query the Reactome Database for specific pathways, select a specific pathway and adjust the volcano plot features (B) Volcano plot with the term ALL used in the Enter Reactome Pathway ID box. This shows all of the experimental dataset (C) Table of pathways generated after putting the term “glucose” in the Enter Pathway Query Term box (D) Volcano plot generated after selecting the Glucose Metabolism pathway and entering the Reactome ID R-HSA-70326 into the Enter Reactome Pathway ID box (E) Table of genes associated with the Glucose Metabolism pathway that are shown in Figure 1D.

### 2.2 Selecting and filtering by pathway

Upon launching the program, the Reactome API is called, and a list of all pathways is generated which in the current version of Reactome (version 90, released October 2, 2024), contain 2742 pathways. The Enter Pathway Query Term box allows the user to enter a term which will then return a list of all of the Reactome Pathways with that term in the name. To view the entire experimental dataset, the term All is input resulting in all of the genes shown as in Figure 1B. When a query term such as “glucose” is input, a table of Reactome pathways containing this term is generated and includes the Reactome ID and the full name of the pathway as in Figure 1C. Next the user can copy and paste the Reactome ID for the selected pathway into the Enter Reactome Pathway box in Figure 1A. Using the Reactome ID for the Glucose Metabolism pathway, R-HSA-70326, the volcano plot shown in Figure 1D is generated which contains only the genes in the experimental dataset that are involved in the Glucose Metabolism pathway. Below the volcano plot is a table of all of the genes in the experimental dataset which are part of this pathway along with the Log2 fold change and adjusted p-value as shown in Figure 1E.

It should be noted that there is a hierarchy in the pathway definitions in the Reactome Database. This is evidenced by the set of pathways returned using the “glucose” query. The top pathway Glucose metabolism is a high level pathway whereas the other three are more specific. Such a hierarchy is found with many other queries as well. For example, the query “metabolism” yields 91 pathways, but includes the high level pathway simply titled Metabolism (R-HSA-1430728). Using this pathway can provide a useful filter while maintaining broad view of metabolism when the working hypothesis includes potential perturbations to metabolism.

### 2.3 Interactive visualization

The plotly package provides several different options to interact with the data. Hovering in the upper right corner of the volcano plot reveals the options to zoom, pan, box select, lasso select, zoom in, zoom out, autoscale and reset axes. The box and lasso select options focus on specific regions of the plot by graying out the genes outside of the selected regions. An option to download a png file of the plot is also provided. Slider bars in the left panel in Figure 1A allow interactive changes of the fold change and p-value thresholds. Note that for simplicity, the p-value threshold is adjusted as the raw value and then translated into the -log10 value on the plot. Gene IDs can be toggled on or off at the bottom and the font size and offset of the labels adjusted by slider bars to optimize the clarity of the plots. A button to download the gene table shown in Figure 1E is at the bottom of the panel.

## 3. Conclusion

The Pathway Volcano tool provides a unique way to simplify the analysis and interpretation of differential expression data by focusing on biological pathways and not just filtering by fold change or statistical significance. This strategy of data reduction helps uncover significant changes that would otherwise be obscured in the crowded region of data near the top of the “cone” of the volcano. The interactivity and the ability to export graphical images and data tables should make this a very helpful tool for researchers working with differential expression data.

## Acknowledgements

Thanks to Dr. Jeffery Baumes and Dr. Roni Choudhury for helpful discussions.

## Funding

This work was supported by NIH/National Cancer Institute SBIR program in collaboration with Kitware Inc.

## Notes

https://github.com/thoconne/PathwayVolcano

### Competing Interest Statement

The authors have declared no competing interest.

